# Therapeutic interventions on human xenografts promote systemic dissemination of oncogenes

**DOI:** 10.1101/2023.04.06.535826

**Authors:** Gorantla V Raghuram, Kavita Pal, Gaurav Sriram, Afzal Khan, Ruchi Joshi, Vishalkumar Jadhav, Sushma Shinde, Alfina Sheikh, Bhagyeshri Rane, Harshada Kangne, Indraneel Mittra

**Affiliations:** Translational Research Laboratory, Tata Memorial Centre, Advanced Centre for Treatment, Research and Education in Cancer, Kharghar, Navi Mumbai – 410210, and Homi Bhabha National Institute, Anushakti Nagar, Mumbai –400094

## Abstract

We generated xenografts of human cancer cells in mice, and using immuno-FISH analysis detected multiple co-localizing signals of human DNA and eight human oncoproteins in brain cells. Signals increased dramatically five days after treatment with chemotherapy, localized radiotherapy or surgery, which could be minimized by concurrent treatment with cell-free chromatin deactivating agents. These results suggest that therapeutic interventions may potentially encourage metastatic spread which is preventable by deactivating cell-free chromatin.

---

Systemic dissemination following successful treatment of the primary tumour remains a common cause of death from the disease. There is mounting evidence that therapeutic interventions themselves may promote metastatic spread**^1^**. This possibility has been raised with respect to all three modalities of cancer treatment viz. chemotherapy**^2,3^**, radiotherapy**^4,5^** and surgery**^6,7^**. Based on our earlier finding that cell-free chromatin particles (cfChPs) that are released from dying cancer cells are potentially oncogenic**^8^**, we hypothesized that therapeutic interventions may disseminate the disease via release of cfChPs from therapy induced dying cancer cells. In keeping with the classical report of Isaiah Fidler**^9^**, we had observed that the vast majority of injected cancer cells had died upon reaching target organs to release cfChPs which integrated into the genomes of target cells. This resulted in dsDNA breaks marked by phosphorylation of H2AX, and activation of inflammatory cytokines NFκB, IL-6, TNFα and IFNγ. Since concurrent activation of DNA damage and inflammation are potent stimuli of oncogenic transformation**^10,11^**, we had hypothesized that cfChPs released from therapy induced dying cancer cells are the critical agents that induce metastatic dissemination of cancer**^8^**.

In the present study, we tested this hypothesis in a pre-clinical model in which we generated xenografts in SCID mice using MDA-MB-231 human breast cancer cells and treated the mice with a single dose of chemotherapy (CT), or localized radiotherapy (RT) to the xenografts or their surgical removal (Sx). The anti-cancer therapies were given either alone or in conjunction with three different cfChPs deactivating agents viz. anti-histone antibody complexed nanoparticles (CNPs)**^12^**, DNase I or a combination of the nutraceuticals Resveratrol and copper (R-Cu).**^13,14^** Combining Resveratrol (R) with copper (Cu) leads to the generation of free radicals**^15^** which can effectively deactivate cfChPs by degrading their DNA components**^13,14^**.

Mice were sacrificed after five days (i.e. on day six) after the above therapies and their brains were harvested and FFPE sections were prepared. The sections were analyzed by immuno-FISH using a human specific whole genomic probe and specific antibodies against human HLA-ABC antigen and eight different human oncogenes viz. c-Myc; c-Raf, p-EGFR, HRAS, p-AKT, FGFR 3, PDGFRA and c-Abl. We detected multiple colocalizing signals of human DNA and HLA-ABC in brain cells of xenograft bearing mice (Fig.1a). Since the HLA-ABC antigen is exclusive to humans and does not exist in mice, this finding provided strong support for the conclusion that DNA fragments carrying the HLA-ABC gene had been released from the dying human xenograft cells and had migrated to mouse brain cells via circulation. We also detected multiple co-localizing signals of human DNA and the eight human onco-proteins that we examined. This finding indicated that the respective oncogenes had also been released from dying xenograft cells and carried via the blood stream to brain cells wherein they had expressed their respective proteins (Fig. 1b).

**Fig.1:**
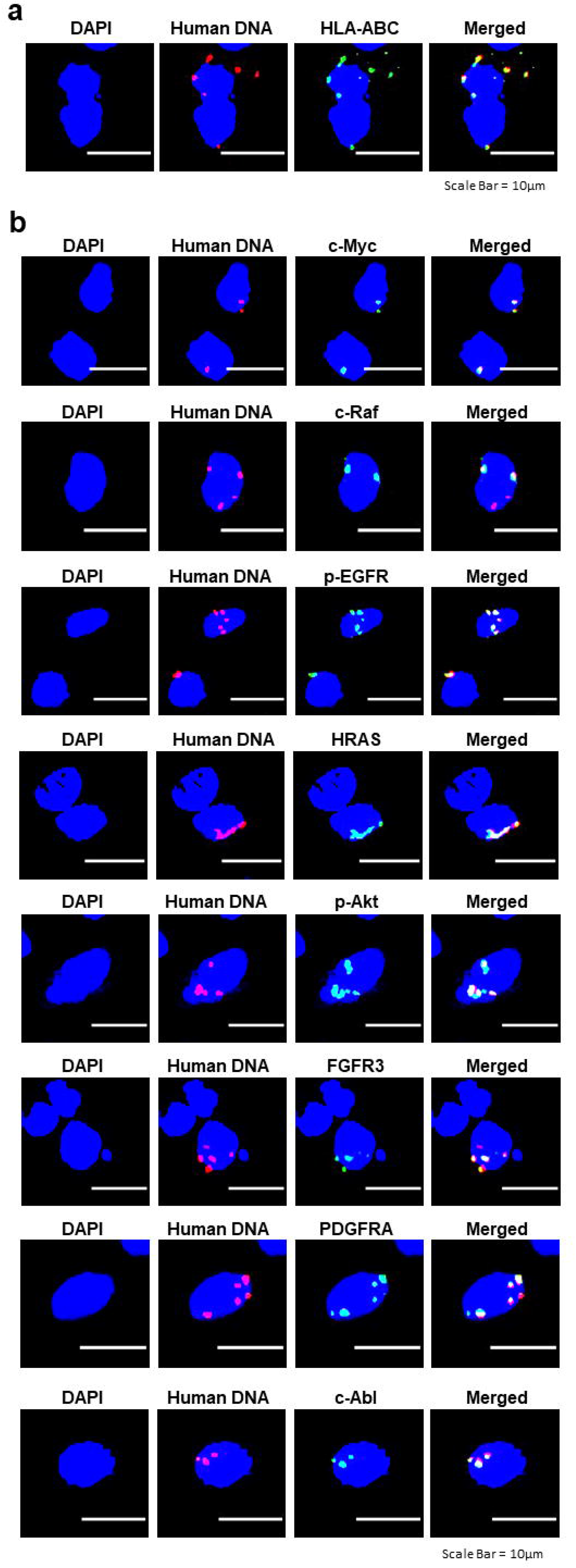
Representative immuno-FISH images of FFPE sections of brains of mice bearing human breast cancer xenograft showing co-localizing signals of human DNA and various human onco-proteins. **a.** Co-localizing signals of human DNA and human HLA-ABC. **b.** Co-localizing signals of human DNA and eight different human onco-proteins.

We next undertook a quantitative comparative analysis of human DNA and c-Myc onco-protein signals in brain cells following the three anti-cancer interventions (Fig. 2, a and b). The analysis was done in a blinded fashion such that the examiner was unaware of the group to which the slides belonged. As expected, no human DNA or c-Myc signals were detected in brain cells of mice in the two control groups viz. control mice without xenograft and control mice without xenograft but receiving anti-cancer interventions (Fig. 2, a and b). However, xenograft bearing mice (without therapeutic interventions) showed that a significant number of brain cell nuclei harbored human DNA (13.73%) and c-Myc onco-protein (3.70%) signals. This data indicated that dying cells of the xenografts had released their DNA into the circulation which had migrated to the brain during the xenograft’s growth span of ~6 weeks. The number of human DNA and c-Myc signals increased markedly following the three therapeutic interventions. With respect to DNA, the maximum increase in signals was seen with CT (13.73% vs 26.33%, p < 0.001), followed by RT (13.73% vs 21.83%, p < 0.01), followed by Sx (13.73% vs 15.83%, p < 0.05). With respect to c-Myc, the number of signals for CT were 3.70% vs 8.08% (p < 0.01) and for RT were 3.70% vs 9.33% (p < 0.001). However, for Sx, the number of signals in untreated and treated groups were not statistically different (3.70% vs 3.37%). Concurrent treatment of mice with all three cfChPs deactivating agents showed remarkable and statistically significant reduction in both human DNA and c-Myc signals with p values ranging between < 0.05 and < 0.01 (Fig. 2, a and b).

**Fig. 2:**
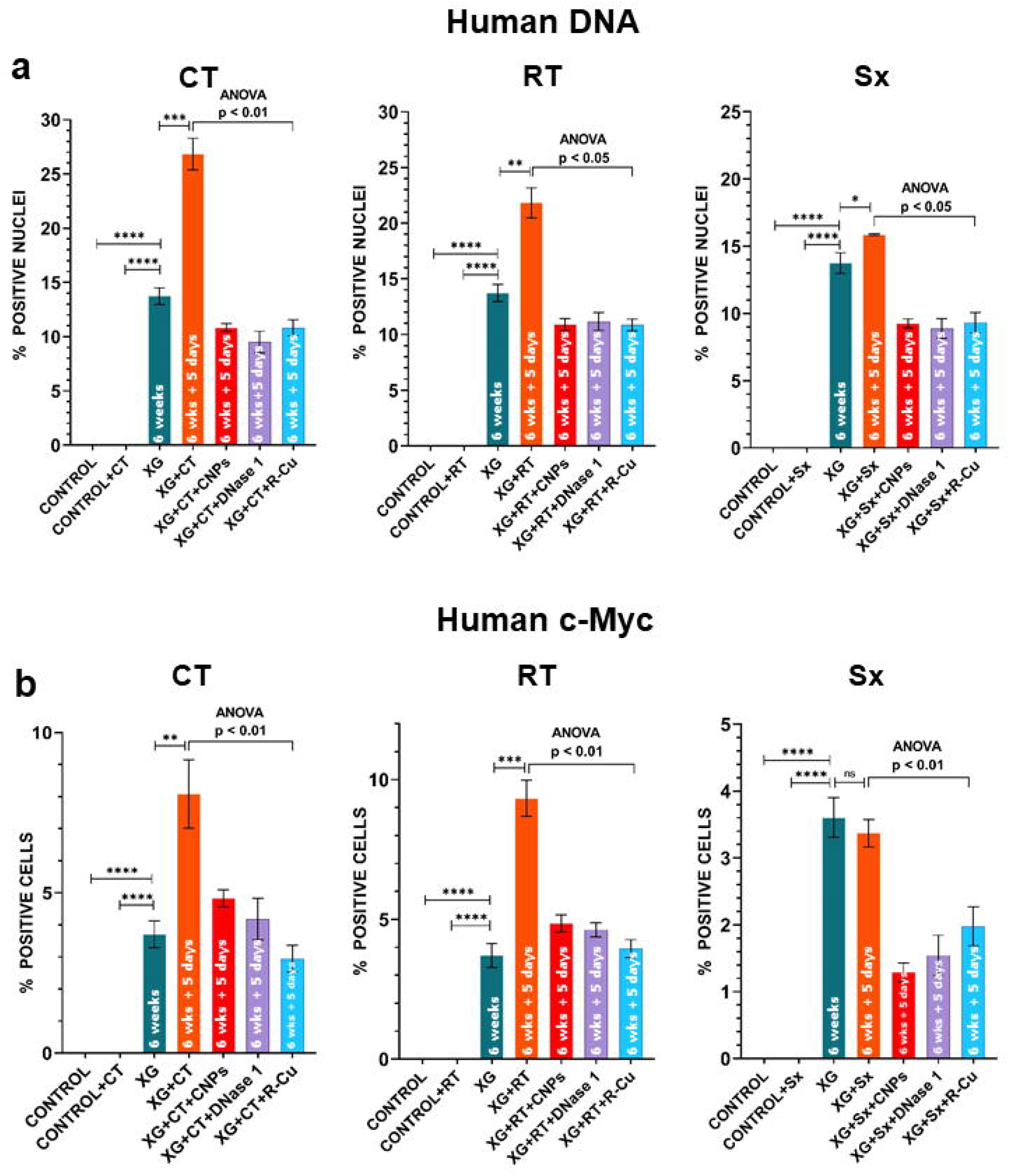
Quantitative analyses depicted as histograms to illustrate that therapeutic interventions promote spread of human DNA and human c-Myc onco-protein to mouse brain cells, and that these can be prevented by concurrent treatment with three different cfChPs deactivating agents. All groups had 4 mice each. **a.** Detection of human DNA by FISH **b.** Detection of human c-Myc onco-protein by immunofluorescence. Statistical comparison between the xenograft bearing group and the two control groups, and that between the xenograft bearing group and the three anti-cancer treatment groups (CT, RT and Sx) was done by two-tailed student t-test. * < 0.05, ** < 0.01, *** <0.001, **** <0.0001. Statistical comparison between the three anti-cancer treatment groups (CT, RT and Sx) and those additionally treated with the three cfChPs deactivating agents was done by One-way ANOVA. * < 0.05, ** < 0.01.

Although systemic dissemination following successful treatment of the primary tumour is a common cause of death from the disease, an explanation for this paradox has remained elusive. We have shown here that DNA fragments released from therapy induced dying cancer cells, and carrying oncogenes with them, can enter into the systemic circulation and be carried to distant organs, such as the brain, with the potential to oncogenically transform their constituent cells. The fact that CNPs, which deactivate cfChPs by binding to their histone components**^12^**, could effectively prevent c-Myc dissemination indicates that the agents that carried the c-Myc oncogene to brain cells were in fact cell-free chromatin particles released from dying xenograft cells. This finding is consistent with our hypothesis that cell-free chromatin particles are the critical oncogenic agents**^8^**.

Whether therapy induced spread of oncogenes to brain cells would lead to development of metastases was not investigated in our short-term study. Nonetheless, the fact that cfChPs have the ability to concurrently activate two critical hallmarks of cancer viz. genomic instability and inflammation**^8^** suggests that prolonged observation is called for. Future long-term experiments should also explore whether cfChPs deactivating agents given concurrently with anti-cancer treatments would prevent metastatic spread. These agents would have the added advantage, when used as adjuncts to cancer treatment, of preventing toxic side effects of chemotherapy**^16,17^** and radiotherapy.**^18^**

## Materials and Methods

### Ethics approval

The experimental protocol of this study was approved by the Institutional Animal Ethics Committee (IAEC) of the Advanced Centre for Treatment, Research and Education in Cancer (ACTREC), Tata Memorial Centre (TMC), Navi Mumbai, India. The experiments were carried out in compliance with the ethical regulations and humane endpoint criteria of IAEC and ARRIVE guidelines.

### SCID mice

Inbred female NOD SCID (NOD.Cg-Prkdcscid/J) mice were used in this study. They were obtained from the Institutional Animal Facility and were maintained according to IAEC standards. They were housed in pathogen-free cages containing husk bedding under 12-h light/dark cycle with free access to water and food. A HVAC system was used to provide controlled room temperature, humidity and air pressure. ACTREC-IAEC maintains respectful treatment, care and use of animals in scientific research. It aims that the use of animals in research contributes to the advancement of knowledge following the ethical and scientific necessity. All scientists and technicians involved in this study have undergone training in ethical handling and management of animals under supervision of FELASA certified attending veterinarian. Animals were euthanised at appropriate time points under CO2 atmosphere by cervical dislocation under supervision of FELASA trained animal facility personnel.

### Creation of xenografts

NOD SCID mice (6-8 week old) were inoculated under the lower dorsal skin with 1 X 10^6^ MDA-MB-213 human breast cancer cells and xenografts were allowed to grow to a size of ~0.125 cm^3^, which usually took ~6 weeks, when the experiments were initiated (Supplementary Fig. 1).

### Study design

The study design is depicted in the schema given below.

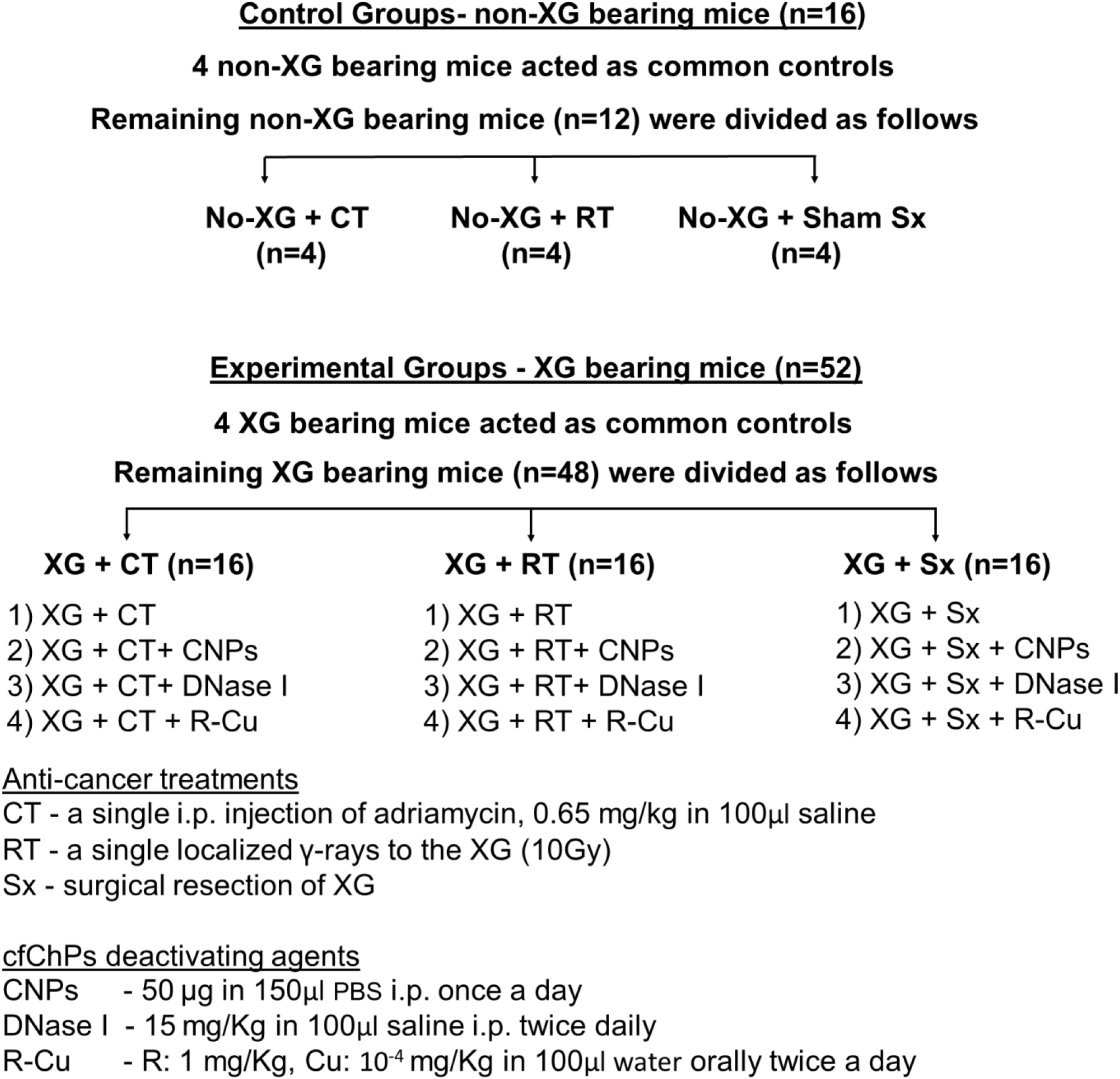

Mice were sacrificed 5 days (i.e. on day 6) after starting anti-cancer treatments; cfChPs deactivating agents were started 4h prior to commensing anti-cancer therapies. Brains of mice were removed, fixed in formalin and FFPE sections were prepared for analysis. <u>Experiment no.1:-</u> Involved XG bearing mice without anti-cancer treatments. Their brain sections were analyzed for detection of human DNA and human onco-proteins by immuno-FISH. <u>Experiment no.2:-</u> Involved XG bearing mice receiving anti-cancer treatments. Statistical comparison was performed on the effects of anti-cancer treatments, with and without cfChPs deactivating agents, on human DNA and c-Myc onco-protein signals in brains cells. Five-hundred cells were analyzed for detection of human DNA signals (at a magnification of x60), and 1000 cells were analysed for c-Myc (at a magnification of x40). Percentage of cell positive for human DNA and c-Myc signals was calculated.

### Details of therapeutic interventions

Therapeutic interventions were initiated when the xenograft has reached the size of ~0.125 cm^3^ which took on an average 6 weeks

#### Chemotherapy

Mice were administered a single i.p. injection of adriamycin, 0.65 mg/kg in 100μl saline

#### Tumour irradiation

Mice were anesthetized using ketamine (80 mg/kg IP) and xylazine (10 mg/kg IP) and placed in a polypropylene box. The box was placed on the couch of a telecobalt machine (Bhabatron-II)^©^ and positioned in such a way that the apex of the tumor was at the center of the aperture of the machine. The rest of the body of the mice was shielded from radiation using 6.5 cm-thick lead shields (Supplementary Fig. 2). The dose delivered to the tumors was 10Gy.

#### Surgical excision

Mice were anesthetized using ketamine (80 mg/kg IP) and xylazine (10 mg/kg IP) and the tumours were surgically removed.

### Details of cfChPs deactivating agents

#### Anti-histone antibody complexed nanoparticles (CNPs)

CNPs were prepared according to the method reported by us earlier**^12^** except that histone H4 IgG was exclusively used for preparing CNPs. CNPs, 50 μg in 150 μl PBS, were administered once a day for 5 days.

#### DNase I

DNase I (Sigma-Aldrich; Catalogue No-DN25-1G) dissolved in saline was administered twice daily at a dose of 15 mg/kg in 100 μl of saline i.p.

#### Resveratrol-copper (R-Cu)

R: 1 mg/Kg in 100μl water, and Cu: 10^-4^ mg/Kg in 100μl water, were administered by oral gavage one after the other twice daily. The sources of R and Cu were (Resveratrol, Trade name—TransMaxTR, Biotivia LLC, USA; Copper, Trade name—Chelated Copper, J.R. Carlson Laboratories Inc. USA).

### Immunofluorescence and FISH analysis

Mice were sacrificed after five days (i.e. on day six) after the anti-cancer therapies, their brains were harvested and FFPE sections were prepared. The sections were analyzed by immuno-FISH using a human specific whole genomic probe and specific antibodies against human HLA-ABC antigen and eight different human oncogenes viz. c-Myc; c-Raf, p-EGFR, HRAS, p-AKT, FGFR 3, PDGFRA and c-Abl. For FISH analysis, 500 cells were examined for detection of human DNA signals (at a magnification of X60) and percentage of cells showing positive signals was calculated. For onco-protein analysis by immunofluorescence, 1000 cells were examined for detection of human c-Myc signals (at a magnification of X40) and percentage of cells showing positive signals for human c-Myc protein was calculated. Details of the FISH probe and various antibodies used in this study are provided in the Supplementary Table 1.

### Statistical analysis

Statistical comparison between the xenograft bearing group and the two control groups, and that between the xenograft bearing group and the three anti-cancer treatment groups (CT, RT and Sx) was done by two-tailed student t-test using <u>GraphPad Prism 6.0</u> (https://www.graphpad.com/, <u>GraphPad Software, Inc., USA)</u>. Statistical comparison between the three treatment groups (CT, RT and Sx) and those additionally treated with the three cfChPs deactivating agents was performed by One-way ANOVA using the same software.

## Acknowledgement

The authors thank Mr. Ashish Pawar for his help in preparing manuscript.

