## Supplementary figures and images for "Therapeutic interventions on human xenografts promote systemic dissemination of oncogenes"

### Supplementary Figure 1

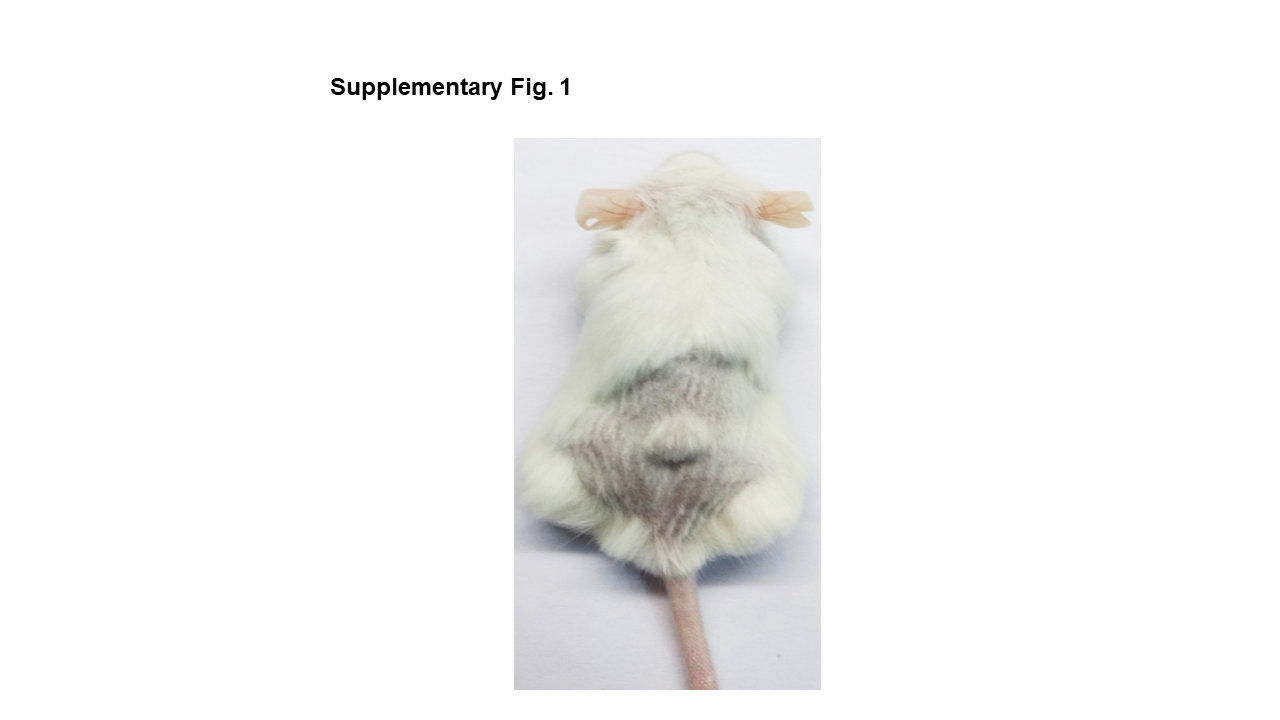

### Supplementary Figure 2

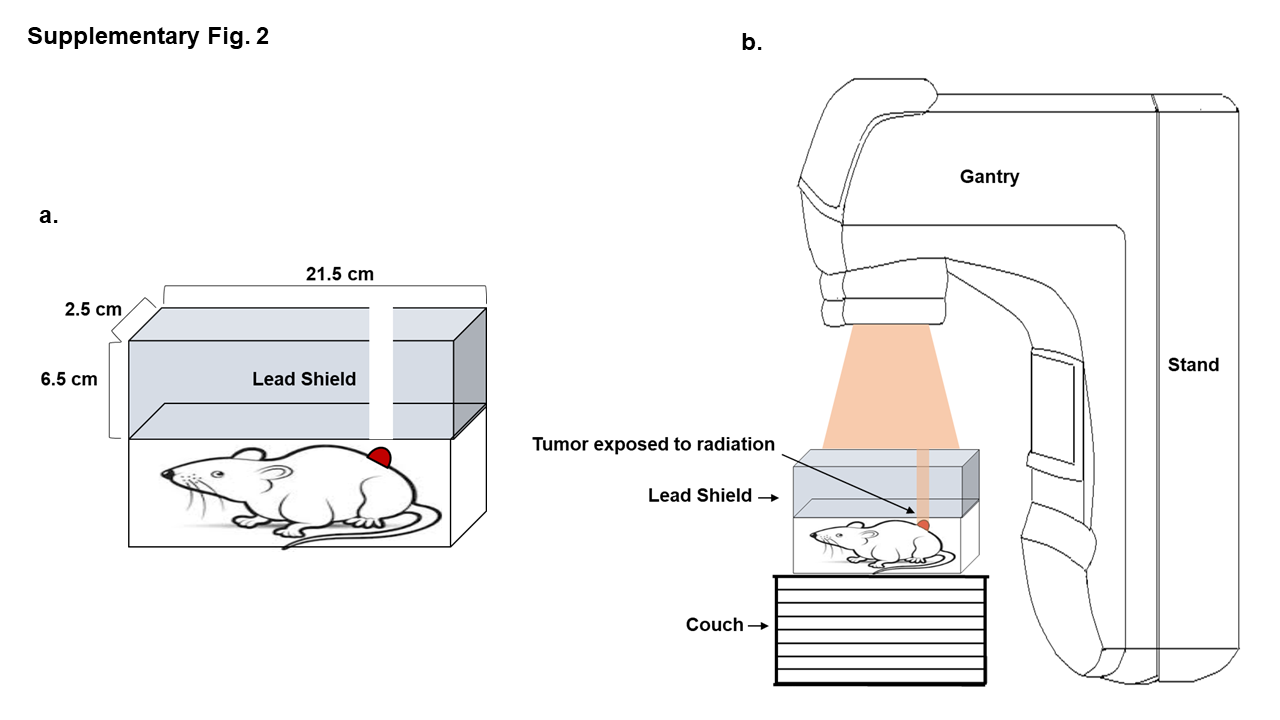
