## Supplementary material for "Therapeutic interventions on human xenografts promote systemic dissemination of oncogenes": Legends to Supplementary Figures

**Supplementary Fig. 1:**

A representative image of a SCID mouse bearing human breast cancer xenograft.

**Supplementary Fig. 2:**

Figure showing set-up for delivering localized radiation to xenografts **(a and b)**.
