## Supplementary Table 1 for "Therapeutic interventions on human xenografts promote systemic dissemination of oncogenes"

**FISH Probe**

| **Sr. No.** | **FISH Probe** | **Catalogue No.** | **Company / Vendor** |
| --- | --- | --- | --- |
| 1. | Human Genomic DNA (probe) Red | Custom synthesized | Applied Spectral Imaging, Israel |

**Primary antibodies**

| **Sr. No.** | **Primary antibody** | **Catalogue No.** | **Company / Vendor** |
| --- | --- | --- | --- |
| 1. | HLA-ABC1 | ab70328 | Abcam, UK |
| 2. | c-MYC | M4439 | Sigma-Merck-Millipore, Germany |
| 3. | c-Raf | MA5-41270 | Invitrogen- Thermo Fisher Scientific, USA |
| 4. | p-EGFR | 4404S | Cell Signaling Technologies,USA |
| 5. | HRAS | Orb38488 | Biorbyt Biotech, USA |
| 6. | p-AKT | 4060S | Cell Signaling Technologies,USA |
| 7. | FGFR 3 | 4574S | Cell Signaling Technologies,USA |
| 8. | PDGFRA | HPA004947 | Atlas Antibodies,USA |
| 9. | c-abl Antibody | PA514762 | Invitrogen- Thermo Fisher scientific, USA |

All antibodies were specific to humans except that for p-AKT which was common for human and mouse.

| **Sr. No.** | **Secondary antibody** | **Catalogue No.** | **Company / Vendor** |
| --- | --- | --- | --- |
| 1. | Goat Anti-Rabbit IgG (H+L) FITC Conjugate Secondary Antibody | AP307F | Merck-Millipore Sigma, Germany |
| 2. | Goat Anti-Mouse IgG H&L (FITC) Pre-Adsorbed Secondary Antibody | ab7064 | Abcam, Cambridge, UK |

**Secondary antibodies**
